# BDNF and Tau as biomarkers of severity in Multiple Sclerosis

**DOI:** 10.1101/236158

**Authors:** Islas-Hernandez Azul, Aguilar-Talamantes Hugo Seacatl, Bertado-Cortes Brenda, Mejia-delCastillo Georgina de Jesus, Carrera-Pineda Raul, Cuevas-Garcia Carlos, Garcia-delaTorre Paola

## Abstract

**Aims:** Determine if serum levels of tau and brain-derived neurotrophic factor (BDNF) can be used as severity biomarkers in multiple sclerosis (MS).

**Patients and methods:** Subjects with MS, older than 18 and younger than 55 years were included; 74 patients with a diagnosis of relapsing-remitting MS (RRMS), 11 with secondary-progressive MS (SPMS), and 88 controls were included. Total tau and BDNF were measured by western blot.

**Results:** Increased tau and decreased BDNF in MS patients compared to controls was found. Total-tau has a peak in RRMS, the second decile of the Multiple Sclerosis Severity Score, and in the lowest Expanded Disability Status Scale and is no different than controls for SPMS patients and the most severe cases of MS.

**Conclusion:** BDNF is a good biomarker for diagnosis of MS but not for severity or progression. Tau appears to have a more active role in the progression of MS.

## Introduction

Neurodegenerative diseases have all been found to have a multifactor etiology. The more we study them, the harder it is to make an association between biomarkers and disease that works for every patient. A simple quantification of biomarkers gives no advantage to the daily clinician. Here, we have studied the biomarkers we thought would give us useful information about patients with multiple sclerosis (MS). A biomarker is defined as a characteristic that is objectively measured and evaluated as an indicator of a normal biological process, pathogenic process, or pharmacologic responses to a therapeutic intervention [1].

For MS, the pathophysiology is expressed in three main compartments: peripheral blood, the blood brain barrier and the central nervous system [2]. Post-mortem cerebral tissue is an excellent source, but its use is impractical when it comes to diagnosis. Hence, corporal fluids such as plasma have been the most common choice for the analysis of biomarkers. Peripheral blood biomarkers are readily available and its perfusion through different organs and tissues can result in the addition or modification of proteins that may vary according to different conditions [2], [3].

Multiple sclerosis is a neurodegenerative, chronic inflammatory, demyelinating disease. The new MS classification divides this disease into relapsing-remitting (RR) and progressive, which is sub-classified in primary progressive (PP) and secondary progressive (SP) [4]. Despite the significant amount of studies in MS, the only validated biomarker is the detection of oligoclonal IgG bands in the CSF, which is used as an adjuvant tool for diagnosis [2].

Individual variations in its clinical course make it hard to evaluate severity in MS. There are three important scales for classifying severity in patients with MS; the most common is the Expanded Disability Status Scale (EDSS), which is a method of quantifying disability and monitoring changes over time [5]. Then there is the Multiple Sclerosis Severity Score (MSSS) that compares individual EDSS scores with the distribution of EDSS in patients who have had MS for the same length of time [6]. Finally, there is the Multiple Sclerosis Functional Composite (MSFC) which is a 3 part standardized, quantitative, assessment instrument developed to reflect the clinical expression of MS [7].

Here, we intended to link a combination of biomarkers, to the severity and the time of diagnosis of MS patients to introduce these tests as an aid for clinicians. We have chosen the brain-derived neurotrophic factor (BDNF) and tau as the biomarkers to be evaluated in serum of MS patients.

BDNF is a neurotrophin that, amongst other things, promotes neuronal survival and potentiates proliferation of oligodendrocytes and myelination. Some authors have proposed a protective role for endogenous BDNF in the nervous system. One study showed increased expression of BDNF receptor TrkB in naked axons, in a murine model of MS, suggesting a regulation of expression by neuroinflammation. In their study, they achieved to diminish the severity of the disease by injecting T-cells that overexpressed BDNF, proposing direct axonal protection by this factor [8].

Furthermore, tau is a microtubule assembly phosphoprotein that participates in axonal locomotion, as well as in intraneuronal transport and was first described as a facilitator for microtubule assembly [9]. Its physiological roles are stabilization of microtubules and maintaining the shape and structural polarization of neurons [10]. Tau has been used as a biomarker for axonal damage [11] since its levels have been reported to increase in various CNS diseases [12].

In contrast with BDNF, reports relating tau and MS are controversial. Some report an increment of the protein in CSF [13],[14], while others a decrement [15] or no significant differences [16]. Terzi and coworkers [14] found increased total tau (T-tau) in patients with MS compared with controls, and revealed a strong correlation between T-tau and the duration of the disease in every patient. Also, Bartosik-Psujek [17] and his group have reported increased T-tau in the serum of patients with MS compared with controls.

We used serum of MS patients to quantify tau and BDNF and used the expanded disability status scale (EDSS), years of diagnosis, and the multiple sclerosis severity score (MSSS) to evaluate the usefulness of these molecules for diagnosis.

## Methods

### Ethics statement

All work was conducted with the formal approval of the human subject committees. The ethical and science committees of the Instituto Mexicano del Seguro Social (study number R-2012-785-049) approved the study. All participants gave written informed consent. This study was performed according to the World Medical Association Declaration of Helsinki. All patients have given written consent for the study, including the acknowledgment of full anonymity. Mandatory health and safety procedures have been complies with in the course of the study.

### Study

This study used a convenience sample, all subjects were recruited between September 2012 and April 2014. Prospective recruitment of all participants was done in the Neurology Service of the Specialties Hospital in Centro Médico Nacional Siglo XXI of the Instituto Mexicano del Seguro Social, Mexico City, Mexico.

We used an observational cause-effect, ambispective study design.

### Subjects

Only subjects that agreed to participate and signed the letter of informed consent were included in this study. Inclusion criteria included subjects with multiple sclerosis diagnoses according to McDonald 2010, older than 18 and younger than 55 years, and any treatment. Exclusion criteria included subjects with any other autoimmune pathology. Controls were also older than 18 and younger than 55 years but with no multiple sclerosis or any other autoimmune pathology diagnosis.

### Consort diagram

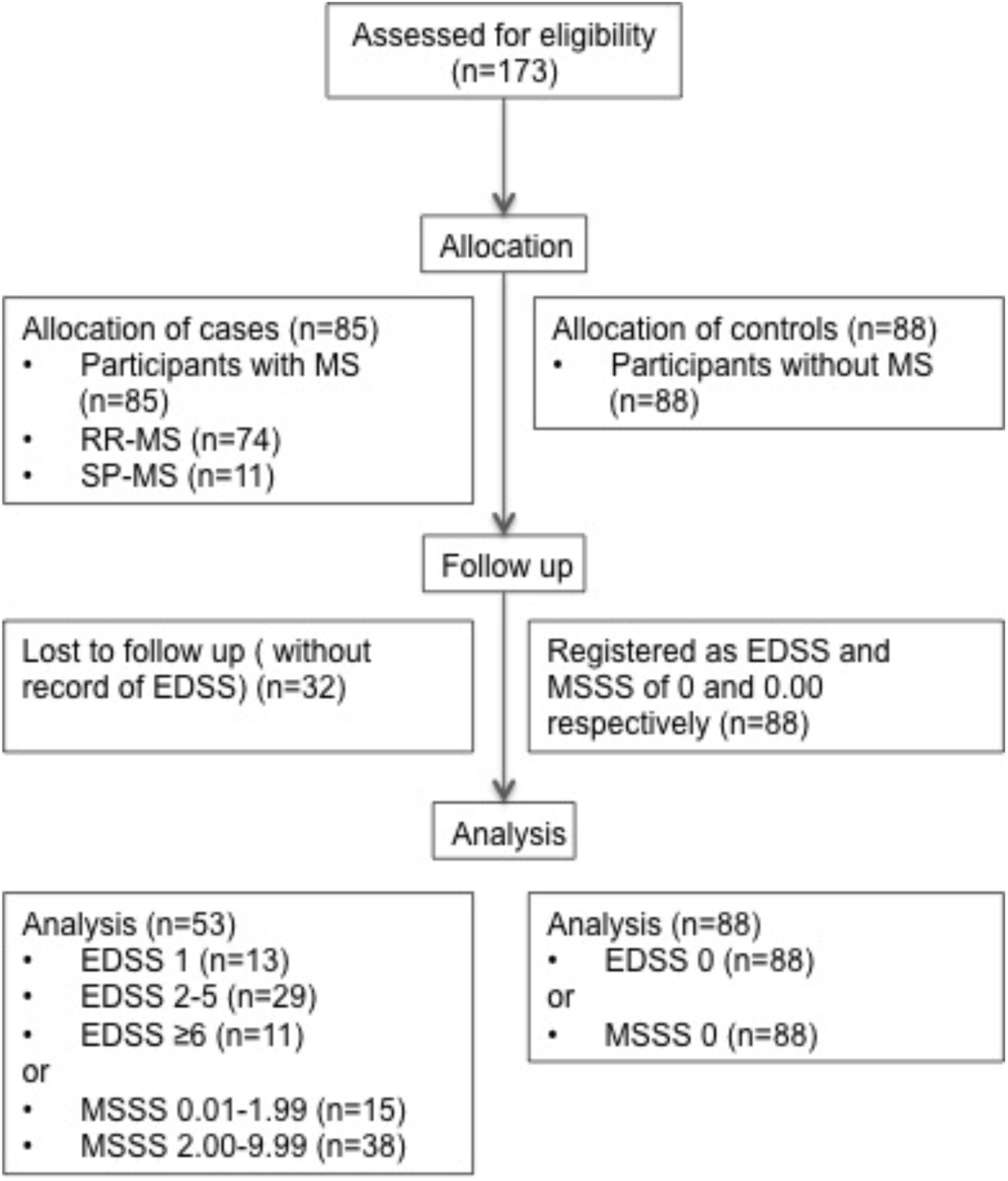

### Measurement of BDNF and Tau

Blood was collected in anticoagulant-free tubes and centrifuged at 2000g for 10min at 4°C. Serum was collected, and samples were stored at ‐20°C for further use. No more than 2 months after the sample was taken passed before they were used. Standard Western blotting analysis was performed to detect BDNF and tau levels in serum. Western blot was selected for this study to increase specificity of detection.

Protein concentrations were measured using the DCTM Protein Assay Kit (BioRad); 60 μg of protein per sample were separated by 14% sodium dodecyl sulphate-polyacrylamide gel electrophoresis. Samples were electrotransferred onto a polyvinylidene fluoride membrane (BioRad) and subsequently blocked with 3% skim milk in TBS-T at 4⍛ overnight. The membranes were incubated 120 min at room temperature with rabbit polyclonal antibody against BDNF (Santa Cruz; sc-546; 1:1000 dilution) or Tau (Thermo Scientific; PA5-16380; 1:1000 dilution) and rabbit monoclonal antibody against Transferrin used as an internal control (Santa Cruz; sc-30159; 1:500 dilution). Blots were then incubated with horseradish peroxidase-conjugated goat anti-rabbit IgG (1:20000; PI-1000, Vector Labs, USA) in TBS-T, for 2 hrs at room temperature.

Every sample was run and analysed twice, controls and MS patients. Protein bands were identified using chemiluminescence (LuminataTM Classico Western HRP Substrate) and measured by densitometry using ImageJ software. We used transferrin as a normalization control; for each sample, relative abundance of the protein of interest was determined by calculating the ratio of the intensity of the signal for the protein of interest to that of the normalization control. Then, an average of both measurements per sample was obtained and used for statistical analysis.

### Statistical analysis

A normal distribution was found for serum values of tau but not for BDNF. BDNF values were analysed with non-parametric tests. Hence, a simple ANOVA was used to evaluate differences between groups of serum tau and a Kruskal-Wallis for BDNF values. The Spearman correlation coefficient was used for BDNF and Tau. A chi-square test was used to compare cut points of BDNF and tau in MS patients and controls. A binary logistic model was used to evaluate the presence or absence of MS with BDNF and tau as independent variables adjusted by the dummy variables of severity by the MSSS as a function of z, as suggested for biomarker’s logistic analysis [18].

## Results

A total of 173 subjects were included in this study of which 102 were women and 71 were men with a mean age of 41±12.0 years; eighty-five were MS patients and eighty-eight controls (without MS). Eighty-six percent of MS patients in this study had an RR diagnosis while 14% had a SP-MS diagnosis. Twenty percent of the MS population was smokers, and 49% had comorbidity (the most frequent being an affective disorder with 12%). While 37% were treated with glatiramer acetate, 63% received other treatment (interferon 12, interferon 6, interferon 8, Fingolimod or Mitoxantrone). The only difference in the measured values was BDNF, specifically in the population with more than ten years of MS diagnostic (24%) with a p=0.1. The main characteristics of the MS patients can be seen in Table 1.

**Table 1.** Clinical data of MS patients

| Table 1. Clinical data of MS patients |  |  |
| --- | --- | --- |
| Variables | Number of patients |  |
|  | RR | SP |
| Patients | 74 | 11 |
| Women | 42 | 8 |
| Men | 32 | 3 |
| 32 patients were lost due to missing data |  |  |
| EDSS |  |  |
| 1 | 13 |  |
| 1.5-5.5 | 29 |  |
| >6 | 11 |  |
| MSSS |  |  |
| 0.01-1.99 | 15 |  |
| 2.00-9.99 | 38 |  |
| RR relapsing remittin, SP secondary progressive, EDSS Expanded disability status scale, MSSS Multiple sclerosis severity score. |  |  |

We found increased Tau and decreased BDNF in the serum of MS patients compared to controls (p<0.05 and p<0.001 respectively). In this context, we found no significant differences for BDNF or Tau in gender of MS patients (p=0.343 and p=0.19 respectively) or controls (p=0.681, p=0.266). Differences between MS patients and controls for BDNF and tau are shown in figure 1.

**Figure 1.**
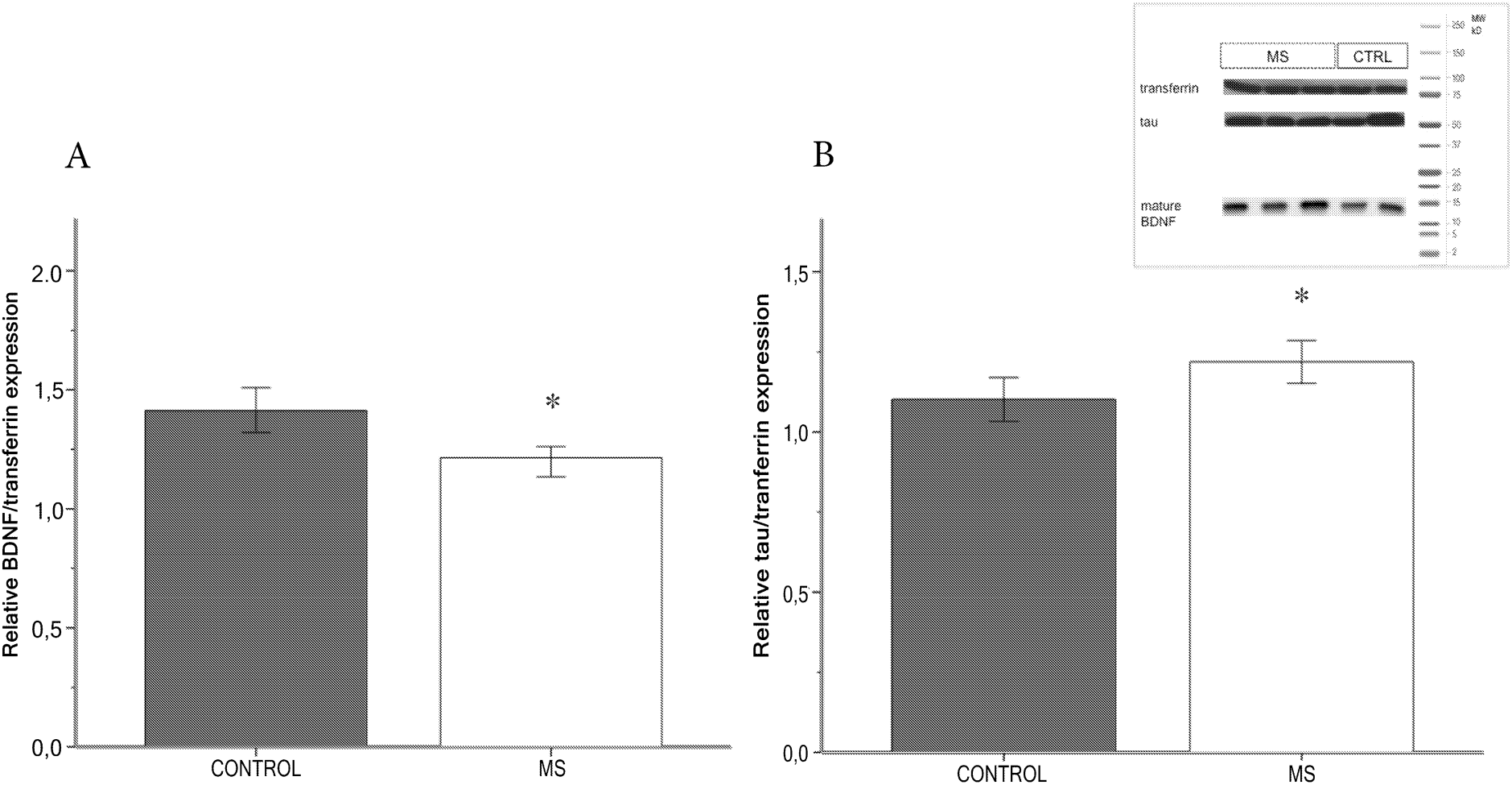
BDNF (1.2148 Median, IQR= 0.4635) and Tau (1.2189 Mean, SE= 0.3111) in the serum of controls and MS patients. A) BDNF serum values are higher in controls (gray) than in multiple sclerosis patients (white). Data is shown as BDNF expression relative to transferrin ^*^ p<0.05 B) Tau serum values in controls (gray) is lower than in multiple sclerosis patients (white). Data is shown as tau protein relative to transferrin expression ^*^ p<0.05. On the top right corner, there is a representative picture of tau, BDNF and transferrin as seen by western blot analysis.

Using a simple ANOVA test, we found differences between controls, RR-MS, and SP-MS patients (p<0.05). A Tukey test showed a significant increment of tau in RR-MS patients as seen in figure 2-A. When the values were compared according to EDSS scores, we found a significant increase in serum tau values for patients with an EDSS of 1 compared with 2-5 and >6 (Figure 2-B).

**Figure 2.**
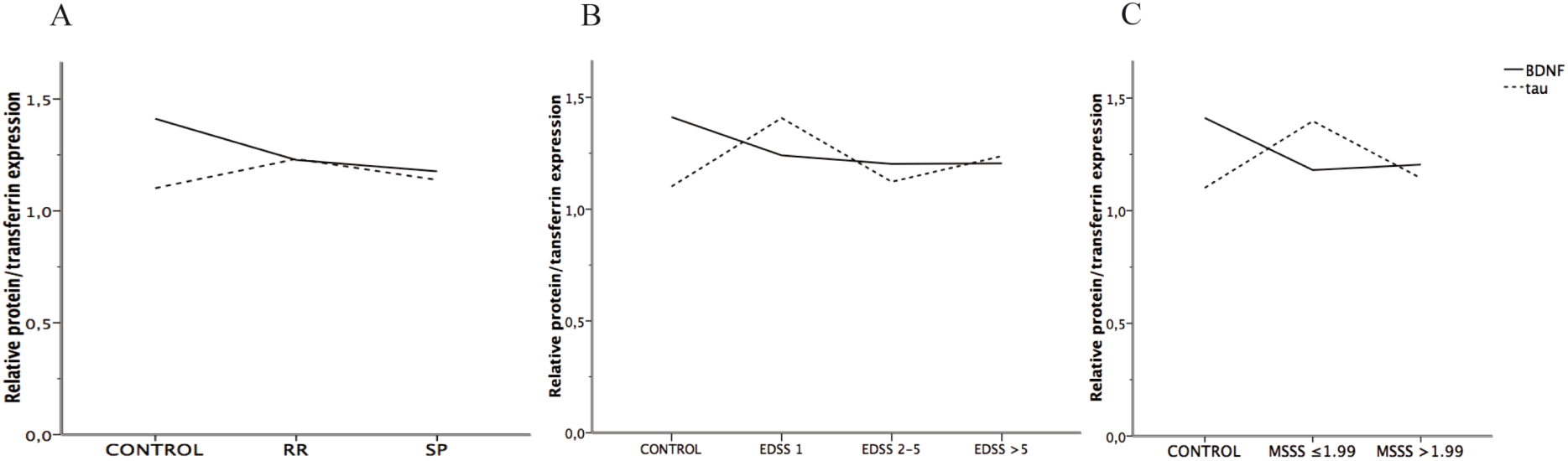
Expression of BDNF and tau relative to severity of multiple sclerosis. A) BDNF (solid line) decreases with both MS clinical courses (RR and SP) while tau (dotted line) increases in RR-MS patients and returns to normal values in SP-MS patients; B) values of BDNF (solid line) and tau (dotted line) according to the EDSS score where BDNF decreases in all categories and tau increases when EDSS is 1 but returns to normal values when EDSS is higher; and C) BDNF (solid line) and tau (dotted line) values according to MSSS, BDNF decreases in MS patients independently of MSSS but tau increases when MSSS is lower than 1.99 and returns to normal values when higher than 1.99.

The same analysis for BDNF serum values with a Kruskal-Wallis test found differences between MS and control groups (p<0.001) with no differences between RR-MS and SP-MS patients. A similar result was found when tested using EDSS values (p<0.05) where differences were only with controls but not between EDSS groups (Figure 2-B).

When the MSSS was used to classify MS patients by severity, a simple ANOVA showed differences for serum tau (p<0.05) with significant differences between the 1^st^ and 2^nd^ decile (0.01-1.99). A Kruskal-Wallis was used to evaluate serum BDNF, a significant difference was found (p<0.05) only in comparison with controls but not between deciles of the MSSS (Figure 2-C).

No statistical correlation was found for BDNF and tau (p=0.153) with a condition index of 1 and a condition number of 7.99. This data showed a low collinearity between variables as a possible non-linear negative association within the participants with a lower MSSS. The cut-off points were established using the descriptive analysis of the data (figure 2). The ANOVA test showed that most of the population with multiple sclerosis was above 1.3oo in the tau/transferrin ratio, and below 1.100 in the BDNF/transferrin ratio. Hence, the values of BDNF <1.1 and tau >1.3 as risk variables were used. A changing association between these biological variables depending on years of diagnosis and MSSS values is shown in Table 2.

**Table 2.** Odds ratio (OR) with a 95% confidence interval (CI) for the association of BDNF, tau and Multiple Sclerosis risk.

|  |  |  |  |  |
| --- | --- | --- | --- | --- |
| **Variables** | **Cut off values** | **P value** | **Correlation (SE)** | **OR (CI 95%)** |
| **MS vs CTRL** |  |  |  |  |
| BDNF | $<1.100$ | *0.005 | 0.214 (0.073) | 2.834 (1.349 – 5.955) |
| tau | $>1.300$ | 0.177 | 0.103 (0.75) | 1.600 (0.806 – 3.178) |
| **Low MSSS vs High MSSS** |  |  |  |  |
| BDNF | $<1.100$ | 0.726 | -0.048 (0.134) | 7.88 (0.208 – 2.989) |
| tau | $>1.300$ | 0.153 | 0.196 (0.142) | 2.45 (0.705 – 8.51) |
| BDNF brain derived neurotrophic factor, MSSS Multiple sclerosis severity score, SE standard error, Low MSSS = 0.01-1.99, High MSSS = 2.00-9.99. P value was obtained using the Chi square test. |  |  |  |  |

Finally, in Table 3, the R^2^ value of the binary logistic model is presented for serum BDNF and tau adjusted by the dummy variables created from the values of MSSS, finding variance values of 73.4-100%.

**Table 3.** Association of BDNF and tau in patients with Multiple Sclerosis adjusted by MSSS.

|  |  |  |  |
| --- | --- | --- | --- |
| Variables | P value | R<sup>2</sup> of Cox and Snell | R<sup>2</sup> of Nagelkerke |
| Complete Model | **<0.01 | 0.734 | 1.000 |
| BDNF | *0.028 |  |  |
| tau | 0.167 |  |  |
| BDNF brain derived neurotrophic value, BDNF cut off value <1.100, tau cut off value >1.300; Low MSSS=0.01-1.99, High MSSS=2.00-9.99, Controls MSSS=0 |  |  |  |

### Summary points

- Our sample allows a confirmation of BDNF as a biomarker of MS diagnosis.
- We have incorporated tau as a biomarker related to function and severity in MS patients.
- Serum T-tau is highly expressed in RRMS patients and when lowest severity is registered.
- Our sample allows associating different levels of severity of the disease as a function of the time of diagnosis with the serum levels of BDNF and tau.

## Discussion

It has been previously suggested that the decreased BDNF levels observed in MS patients are due to a deficient regulation of the immune response, either by immune cells exhaustion due to attempts at tissue regeneration and repair of neuronal damage, or by the inhibition of its neuroprotective function [19], [20]. In agreement with other studies [19],[8], we found serum BDNF decreased in MS patients compared with controls.

When the MSSS was applied to our population [6], we found no differences by severity, but a decrement in comparison to controls. Therefore, BDNF seems like a good biomarker for diagnosis since it has always been found to decrease in MS patients, but not for severity or progression, at least not for our population.

Recently, a decrement in plasma BDNF in male patients newly detected with MS was found [21], and it was proposed that this early decrease is due to the stress of being diagnosed with the disease. On that matter, relapse of MS patients has also been associated with anxiety and depression episodes. Furthermore, MS patients can debut with depressive symptoms and resistant to treatment [22].

On this same lane of though, leading epidemiological studies of medical comorbidities in patients with MS show major depressive disorder (MDD) as the most common, reaching up to 7 fold in MS population relative to controls [23]. One of the difficulties of this study was the fact that we did not take comorbidity into account because of the lack of information, especially those related to mental health. MDD would be of particular interest, since these patients, even when MS is not present, have also shown decreased serum BDNF levels [24], [25].

We found a low correlation of BDNF with MS diagnosis; values under 1.1 were 1.8 times more associated with MS than with controls. It appears as if BDNF decreases at the beginning of the disease, where one of the most common clinical manifestations is chronic or recurrent depression, but acute affective symptoms are not always expressed. Accordingly, suicide has been found to occur up to 3 times more in MS patients than in the general population, becoming one of the leading causes of death in this particular group [26].

Also, interleukins 1a, 1b, and 6, TNF-alfa, or C-reactive protein have been widely described elevated in MS patients [27] as in patients with psychiatric manifestations mainly expressing psychosomatic symptoms [28],[29]. At this time, we were unable to measure inflammation biomarkers, but we believe that the difference found between RR and SP could be related to the inflammation process of MS, where RR-MS patients have been found to have a more acute inflammation response than SP-MS patients.

We also evaluated T-tau in the serum of MS patients, and unlike BDNF, tau has been reported to increase [13],[14],[30], diminish [15] or even have no change [16] in patients with MS. For our population, we found elevated serum T-tau in MS patients compared with controls in concordance with some of the literature [13],[14],[30].

For tau, there seems to be a different dynamic than for BDNF, in which the latter decreases with the disease but has no changes within the progression. Tau, on the other hand, is increased in RR-MS, in the second decile of the MSSS and even in the lowest EDSS, which is what we expected. However, in SP-MS and the most severe cases measured by either EDSS or MSSS, tau returns to normal levels, suggesting an active role for this protein in MS.

On that matter, Jaworski and coworkers found a reduction in CSF T-tau between RR and SP-MS patients, proposing that since the SP stage has a predominant neurodegenerative mechanism, that results in a decreased neuronal density and higher cerebral atrophy, that ends in a loss of tau resources and thus lower T-tau in CSF. The negative correlation between tau concentration in CSF and the degree of motor disability measured by EDSS proposed that tau may be indirectly reflecting the extent of neuronal loss [15].

Nonetheless, our measurements were done in serum and not in CSF, and it has been stated that CSF and serum tau have no correlation [31]. On this matter, it has been suggested that tau in serum is found mainly when neurologic disorders are associated to damage of the brain-blood barrier (BBB) [31]. This has been the case for strokes or head traumas, when indeed the BBB is thought to be impaired rapidly, leading to quick leakage of large concentrations of tau protein into the CSF wile on chronic conditions, tau is continuously released into the CSF. The current theory about MS pathophysiology proposes that during relapses, the protein is released from damaged axons, but its levels are limited because of axon and BBB repair during recovery [15],[32].

A malfunction of the BBB is one of the earliest cerebrovascular abnormalities seen in MS; the breakdown is thought to be transient although recurrence may be observed and this is believed to be in part a cause of the disease [33]. Proinflammatory cytokines promote BBB breakage when a relapse occurs in MS patients. Hence, we believe our results have more to do with Sjörgen's hypothesis of tau being a BBB damage marker in serum [31] instead of an extrapolation of what Jaworski suggested from his results of T-tau in CSF of a decreased neuronal density in SPMS patients [17].

The truth is, not enough has been studied when it comes to serum tau, in part because of the lack of standardized methods to measure it due to its low levels [34], and also because of the interpretation of its presence in serum [15].

The use of BDNF or tau as biomarkers for MS is not new. We attempted not only to identify differences between patients and controls or even differences between RR and SP, which we did find. Our purpose was also to evaluate the usage of these biomarkers for clinicians. So, even though significant changes were observed for both selected biomarkers, when we approached the data with a more careful statistical study, we found no use for progression diagnosis.

It does not mean that BDNF or tau have no biological meaning or clinical use. A proven effect of BDNF is directly related to remyelination [35], and neuroprotection [8] in neuroinflammation and the changes in this neurotrophic factor can help with individual assessment of the patient. We believe particular attention in regards to levels of BDNF and MS comorbidity with MDD is warranted. And as for tau, it can have application as a possible prognostic factor to predict progression for CIS to MS as it has been shown in some initial studies [32] but not as a single prognosis biomarker.

Henceforth, in order to use BDNF and tau as biological variables in a predictive model of MS, we believe a few points must be considered: 1) for our population the cut point for BDNF was <1.1 and for tau >1.3 to be used as associated factors to MS; 2) tau does not have a linear association with MSSS with an increase in the patients grouped in the second decile and then returning to normal values, 3) establish the absence of psychiatric comorbidities (mainly affective and anxiety-related) or at least introduce inflammatory variables as a confounding factor, and finally, 4) incorporate image criteria as an independent covariate.

## Conclusion

BDNF seems like a good biomarker for diagnosis since it has always been found to decrease in MS patients, but not for severity or progression, at least not for our population. Tau was increased in RRMS, in the second decile of the MSSS and even in the lowest EDSS. We also found that in SPMS and the most severe cases measured by either EDSS or MSSS, tau returns to normal levels, suggesting an active role for this protein in MS.

## Appendices

Table 3. Association of BDNF and tau in patients with Multiple Sclerosis adjusted by MSSS. The adjustment for the dummy clinical variables (MSSS) corrects the association of BDNF and tau with the presence o Multiple Sclerosis since the association between the biological and clinical variables is non-linear.

